# On the contrast response function of adapted neural populations

**DOI:** 10.1101/2023.10.06.561309

**Authors:** Elaine Tring, Mario Dipoppa, Dario L. Ringach

## Abstract

The magnitude of neural responses in sensory cortex depends on the intensity of a stimulus and its probability of being observed within the environment. How these two variables combine to influence the overall response of cortical populations remains unknown. Here we show that, in primary visual cortex, the vector magnitude of the population response is described by a separable power-law that factors the intensity of a stimulus and its probability.

## Main

The responses of neurons in primary visual cortex to drifting sinusoidal gratings depend on their intensity, or luminance-contrast. The resulting contrast-response curves are often fit by the Naka-Rushton equation^1,2^, which describes a sigmoidal shape with a restricted linear range and saturation at low and high contrasts^2-5^. Here, we focus on the vector magnitude of the population response^6^, defined as the Euclidean norm of the population vector r = (*r*_1_, *r*_2_,…,*r*_*n*_), which we denote by *r* = ‖*r*‖. As the shape of contrast response curves vary substantially from one neuron to the next^7^, and some neurons in mouse V1 are suppressed by contrast^8,9^, it is not immediately clear how *r* should depend on stimulus contrast. Here, we offer the first experimental measurements of such dependence.

Besides stimulus contrast, the response of cortical neurons is affected by the probability that a stimulus is encountered in the environment through adaptation^5,10-17^. Indeed, we have previously reported that for a stimulus set of fixed, high contrast, the magnitude of the population response is linked to the probability that a stimulus is observed within the environment via a power law^18^, *r* ∼ *p*^*β*^ (on the condition that *r* is approximately constant for a uniform distribution on the stimulus set). The exponent *β* is negative, implying that the magnitude of the population response decreases as the probability of a stimulus increases.

Our goal in the present study is to investigate how the vector-magnitude of a population response depends jointly on stimulus contrast and probability. In other words, how does adaptation change the contrast response function of neural populations? To anticipate the results, we extend our previous findings by showing that response magnitude as a function of stimulus probability and contrast is described by a separable power law, *r* ∼ *p*^*β*^*c*^*δ*^. Here, *r* represents the magnitude of the response, *c* denotes stimulus contrast, *p* is the stimulus probability, and the exponents satisfy *β* < 0 and *δ* > 0.

We obtained this result by recording from excitatory neurons layers 2/3 in mouse primary visual cortex (area V1) using *in-vivo*, two-photon imaging (**Methods**). We measured neural responses to the presentation of a sequence of rapidly flashed, sinusoidal gratings with orientations drawn from von Mises distributions representing three, different statistical environments, with peaks centered at 0, 60, and 120 deg (the concentration of the distribution, *κ* = 1.2, was constant across environments) (**Fig 1**). For each orientation, the contrast was drawn uniformly from a set of 7 values logarithmically spaced between 5% and 100%. The data were analyzed by computing, for each environment, the average magnitude of the population response as a function of contrast and orientation. We then averaged the responses across environments after aligning them to the peak of the orientation distribution. This step relies on the distributions across environments being the same except for the location of their peaks (we will later describe how to relax this condition). Such averaging minimizes variability due to tuning inhomogeneities in a population, where the norm of the population depends on stimulus orientation when the probability distribution is uniform^19^. Finally, each orientation can be associated with its probability of appearance. The resulting dataset can be interpreted as describing how the magnitude of the population response changes as a function of stimulus strength (its contrast) and stimulus probability, *r*(*c, p*).

**Figure 1.**
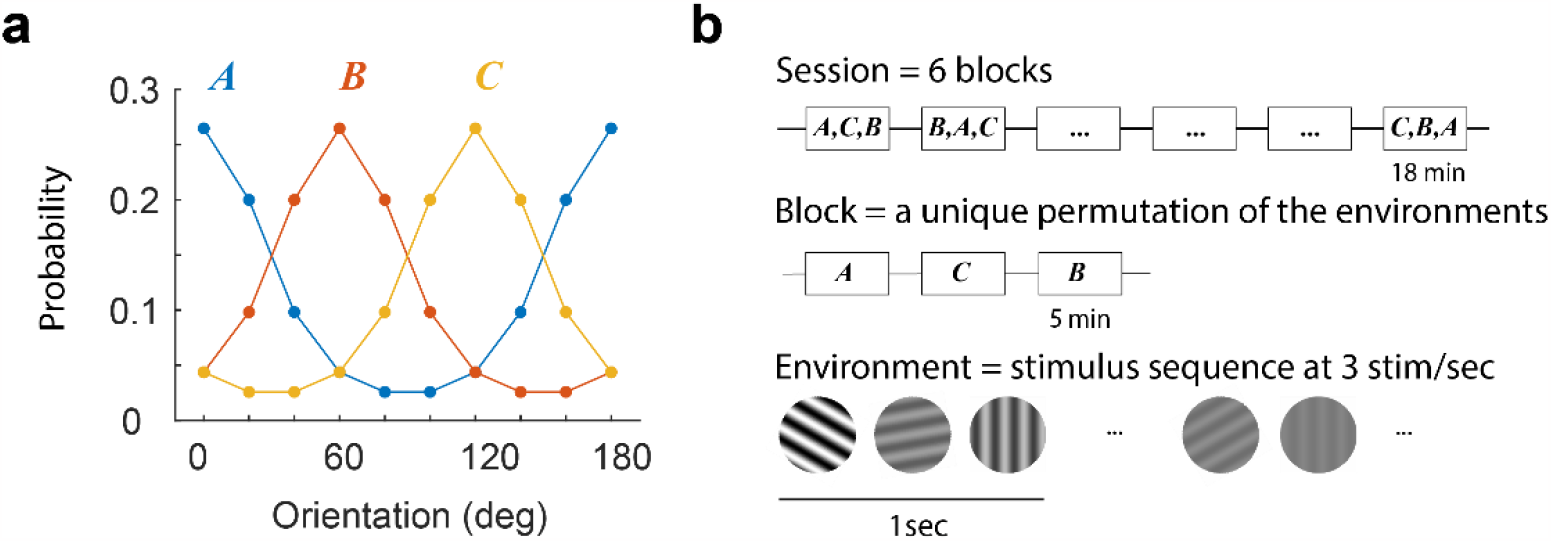
Experimental protocol. (a) Sessions included the presentation of three environments, *A, B*, and *C*, associated with von Mises distribution centered at 0, 60, and 120 deg, respectively. These distributions determine the probability that a stimulus is presented within each environment. The contrast of any selected grating was drawn uniformly from a set of 7 contrast levels equally spaced on a logarithmic scale from 5% to 100%. **b**, A session consisted of six blocks, each containing a unique permutation of all three environments. Each environment was presented for 5 min. Within an environment, stimuli were shown at a rate of 3 per second. A blank screen was presented for 1 min between environments. From one session to the next, the order of the permutations was randomized. Stimuli were presented through a 20 deg circular window centered on the aggregate receptive field of the population (see **Methods**).

The structure of *r*(*c, p*)can be visualized as a 2D pseudo-color image (**Fig 2a**). Each row in the image describes how the magnitude of the response changes with contrast. Each column describes how the response changes with the probability of the stimulus (the peak of the orientation distribution is centered at 0 deg and the data averaged across environments). As expected, the magnitude of population response decreases with the increasing probability of a stimulus (adaptation), and the response increases with contrast. How can the surface *r*(*c, p*)be described in terms *c* and *p*? We discovered that a three-parameter model of the form *r* = *A p*^*β*^*c*^*δ*^ fits the data well. To fit the model, we first take logarithms, resulting in a linear model, log *r* = log *A* + *β* log *p* + *δ* log *c*, with high quality fits (*R*^2^ = 0.95 ± 0.02, mean ± 1SD, **Fig 2b,c**). The coefficient β is negative (−0.34 ± 0.07, mean ± 1SD, *n* = 14), meaning that the higher the probability of a stimulus, the lower the response (summary statistics in **Supplementary Table 1**). The coefficient *δ* is positive (+0.59 ± 0.10, mean ± 1SD, *n* = 14), meaning that the higher the contrast of a stimulus, the higher the response. As a result of the separability of this equation, stimulus contrast and probability trade off against each other in a simple way – a two-fold increase in the probability of a stimulus can be countered by increasing its contrast by a factor of 2^−(*β*/*δ*)^≈ 3.33 leaving the response magnitude invariant. One would be justified in saying that the log-probability of a stimulus and its log-contrast trade in the same “currency.”

**Figure 2.**
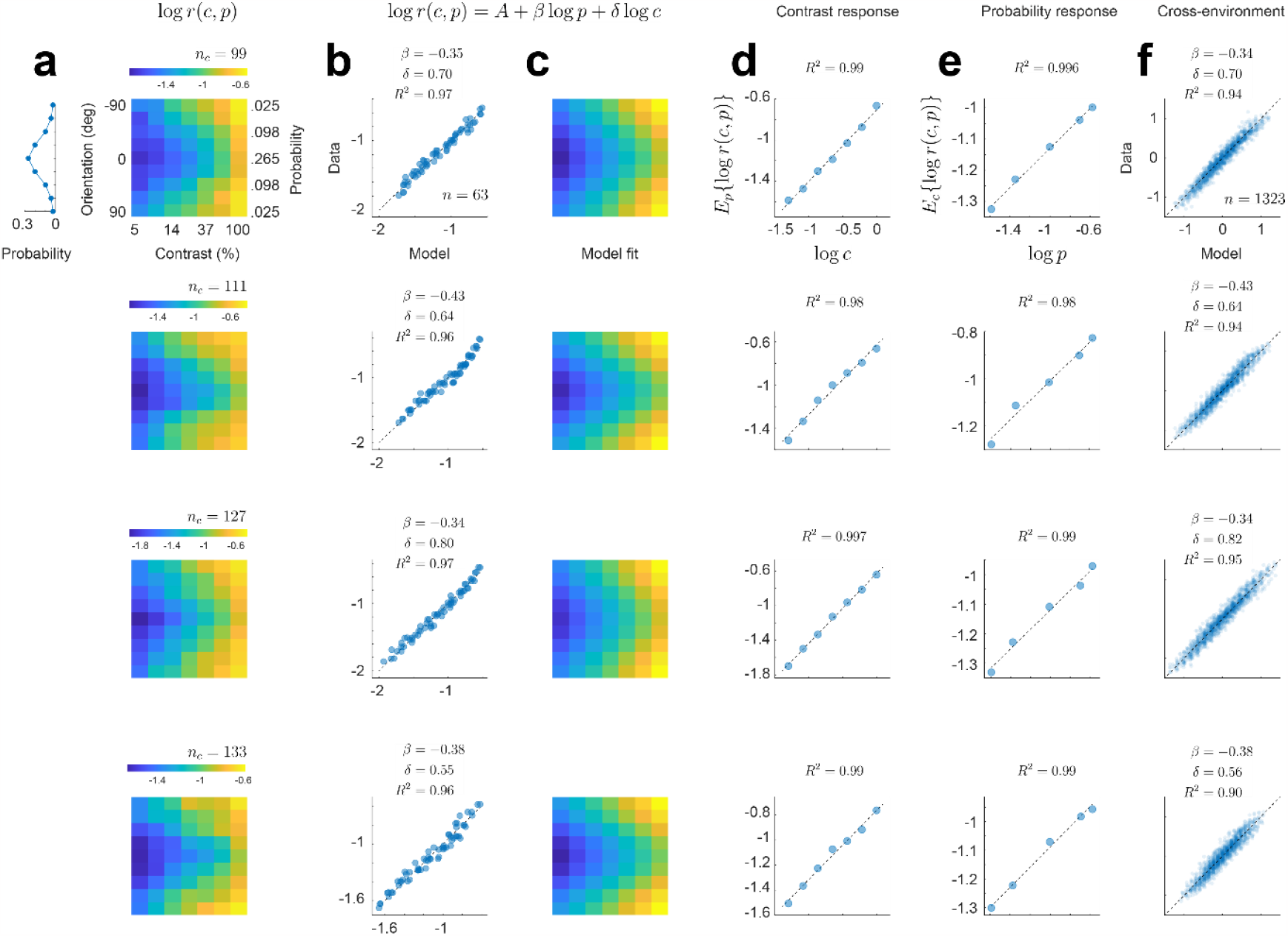
Cortical populations combine stimulus probability and intensity via a power law. Each row of panels describes the outcome of a different experimental session. Unless noted otherwise, the labels and ranges of the axes on the top row apply to all others. **a**. Empirical dependence of response magnitude with stimulus probability and contrast. The pseudo-color image represents the vector-magnitude of the population in a logarithmic scale (scalebar). The number of cells selected is shown by the value of *nc*. The contrast of stimuli varies along the *x*-axis, while the probability (or orientation) varies along the *y*-axis. The results of the three environments were averaged by aligning the peak of the probability distribution with 0 deg. The distribution of orientations (after alignment) is shown by the graph on the left. **b**. Fits of the model *r* = *A p*^*β*^*c*^*δ*^ to the data. The x-axis represents the fits of the model and the y-axis the data. The exponents estimated appear at the inset. The goodness of fit, *R*^2^, is higher than 0.95 in all cases, indicating the quality of the model. **c**. The fits shown in **b** now represented as in the same format as the data in **a. d**. Average contrast response of the population obtained by averaging the rows of the images in **a**. The data falls approximately on a line in log-log coordinates. **e**. Average probability response of the population obtained by averaging the columns of the images in **a**. The data falls approximately on a line in log-log coordinates. **f**. Modeling the ratio population responses across environments as the product between a power of the ratio of the stimulus probabilities and a power of the ratio of the stimulus contrasts. The data are well fit by a line in log-log coordinates. The estimated exponents and goodness of fit appear at the inset. In these and all subsequent analyses the logarithm is base 10.

A more general method of analysis that avoids averaging the data across environments consists of modeling how the ratio of responses between environments changes as a function of the ratio of contrasts and the ratio of probabilities for each stimulus. Given an environment *X*, we denote by *r*(*c*_*X*_, *p*_*X*_(*θ*))the average response to a grating with orientation *θ*, contrast *c*_*X*_, and probability *p*_*X*_(*θ*). Then, our analyses show that the ratios of response magnitudes between two environments, *X* and *Y*, can be described by,

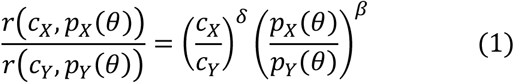

Taking logarithms results in a linear model we can fit to estimate the values for the exponents. Scatterplots between the measured ratios (taking pairwise combinations of the three environments and all stimulus orientations) and their linear fits in log-log coordinates illustrate the high quality of such description (**Fig 2f**).

It is reassuring that the estimates of *β* and *δ* obtained by fitting the data averaged across environments (*β*_*a*_ and *δ*_*a*_) or by fitting the ratios across environments (*β*_*r*_ and *δ*_*r*_) are nearly identical (**Fig 3a**). Moreover, the exponents *β* and *δ* are significantly anti-correlated and well fit by the relationship, *δ* ≈ −1.72 *β* (*ρ* = −0.65, *p* = 0.011)(**Fig 3b**). This correlation means that the more “linear” the population contrast response function of a population is (that is, the closer the value of *δ* is +1), the stronger the population adapts (its exponent *β* is closer to perfect adaptation, attained for *β* = −1). The relationship linking ratios across environments (Equation 1), is rather general and also applies in scenarios where the shape of the distribution differs from one environment to the next (**Fig 3c**).

**Figure 3.**
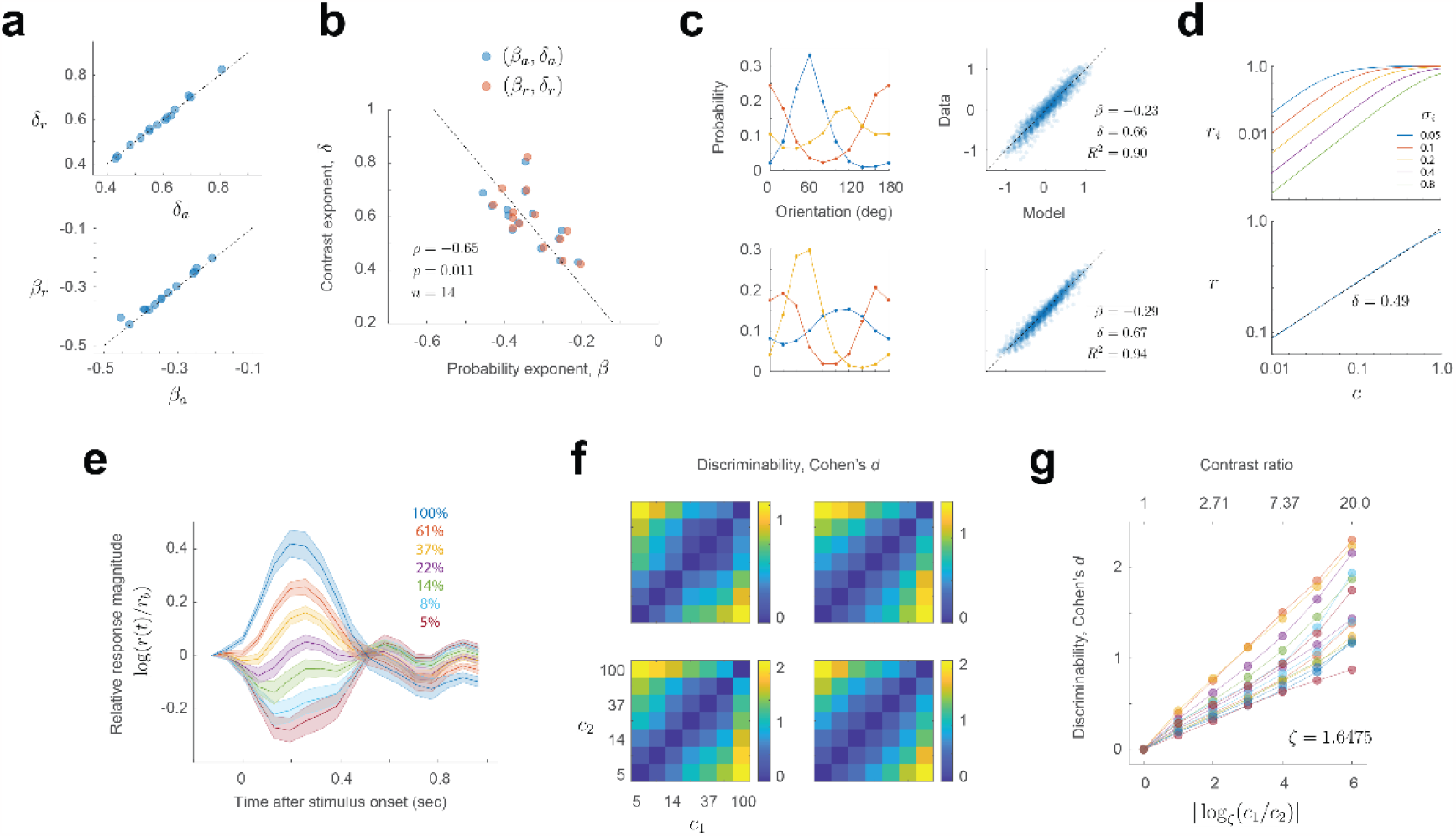
Additional properties of the power law. **a**. Estimates of the exponents agree for the two methods employed: averaging across environments or modeling the ratio of responses across environments. The dashed line is the identity line. **b**. There is a significant, negative correlation between the exponents. Dashed line is the best linear fit. **c**. Modeling the ratio of population magnitudes between environments via a power law holds when the stimulus distributions differ across environments. Each row of panels shows the result from different experiments. **d**. A simple model incorporating a wide distribution of semi-saturation constants in a population of cells with Naka-Rushton contrast responses can produce an almost linear relationship between magnitude and contrast in log-log coordinates. *Top*, contrast response functions of a sample of cells with different semi-saturation constants. Note the graph is in double logarithmic scale. *Bottom*, population contrast response function of a population where the semi-saturation constant is drawn uniformly between 0 and 1. Blue line represents the theoretical model. Dashed, black line is the best linear fit. **e**. Dynamics of population responses as a function of stimulus contrast. The responses to a contrast in each trial are normalized by the activity just preceding stimulus onset, *r*_0_. The responses are averaged across all experimental sessions (*n* = 14). Shaded regions represent ± 2 SEM. Solid lines represent the mean. **f**. Discriminability (Cohen’s *d*) between gratings of the same orientation at two contrasts levels (*c*_1_ and *c*_2_). **g**. Average discriminability is approximately linear with the logarithm of the ratio between the contrasts to be discriminated. The set of contrasts used are spaced logarithmically from 5 to 100% in 6 steps. The ratio between adjacent contrasts is ζ = (100/5)^(1/6)^≈ 1.6475. Thus, we found convenient to express the logarithm represented along the *x*-axis in base ζ.

We conclude that a simple model, log *r* = log *A* + *β* log *p* + *δ* log *c*, accounts for the average magnitude of population responses as a function of stimulus contrast and probability (**Fig 2a-c**). The data showed no evidence of an interaction between the two terms, allowing stimulus contrast and probability to trade off against each other. The model implies that, in any given state of adaptation, the population contrast response function is just a line in log-log coordinates, which does not saturate within the range of contrasts tested. Adaptation at the level of single cells can often be described as a shift in the semi-saturation constant. At the population level, the description is even simpler – changes in adaptation merely translate the contrast response function vertically in log-log coordinates (by the term *β* log *p*). A more general model (Equation 1) accounts for the data without the need of averaging across environments (**Fig 2f**).

At first, one may be puzzled that the population contrast response curve does not saturate. How is this possible if the contrast response of individual neurons, as described by the Naka-Rushton equation, show saturation at low and high contrasts. One possible explanation is that this result is due to a large spread in the distribution of semi-saturation constants across the population^20^. To see how such heterogeneity could shape the contrast response of the population, consider a simple “toy model” where the *i* − *th* cell in the population has a contrast response described by 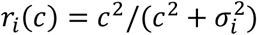. Let us also assume that the semi-saturation constant σ_*i*_ is uniformly distributed between 0 and 1 (here, the value 1 represents 100% contrast), and the exponent is fixed at 2 (**Fig 3d, top**). Then, the vector magnitude of the population as a function of contrast can be shown to be, 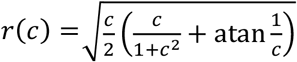. This function turns out to be nearly linear in log-log coordinates (with an exponent *δ* ≈ 1/2), replicating a key feature of our data (**Fig 3d, bottom**).

Further insight into this phenomenon emerges by making an analogy between this model and an audio level meter LED display, where a row of LEDs progressively lights up as the level of the audio signal is increased. If each LED is thought of as a “neuron” that is firing or not, so the contrast response function is a simple step from zero to one that occurs at some level, the row of LEDs is analogous to a population of neurons with semi-saturation constants distributed uniformly. The number of LEDs that are turned on increases linearly with the intensity of the signal and, therefore, the “population norm” increases with the square root of the signal intensity – in other words, the LED meter satisfies a power law with *δ* ≈ 1/2. These models illustrate how heterogeneity in the responses of cortical neurons may be required to endow the population with desirable properties, such as a linear contrast response function in log-log coordinates. An additional feature of these experiments is that the visual presentation protocol, intended to mimic the changes of the retinal image during saccadic eye movements^15,21-23^, keeps the mean population activity at a moderate baseline. The response to a high contrast stimulus drives the responses above this level, while low contrast stimuli drive the responses below the baseline (**Fig 3e**). The ability of the network to modulate its activity above and below a moderate baseline may also help the population to avoid saturation at low and high contrasts.

Finally, we examined how the discriminability between two gratings (of equal orientation) depends on their contrasts *c*_1_ and *c*_2_ (**Fig 3f,g**). We found that discriminability, computed as Cohen’s *d* is proportional to the logarithm of *c*_1_/*c*_2_ (it is convenient to take the base of the logarithm equal to the ratio between adjacent contrasts in our experiments, ζ = 1.6575) (see **Methods**). Each of our experiments generated an approximate relationship *d* ∼ *k* |log_ζ_ *c*_1_/*c*_2_| (**Fig 3g**), but the slope *k* varied between experiments (mean 0.25 ± 0.07, mean ± 1SD). This result is puzzling as it implies the population obeys a neurometric Weber’s law when, in humans, contrast discrimination thresholds violate it^24^. Unfortunately, to our knowledge, there are no measurements of behavioral, suprathreshold contrast discrimination in the mouse to compare our data to at present.

Altogether, our findings reveal that a simple relationship accounts for the interplay between stimulus strength and probability in driving the magnitude of population responses. To verify that non-linearities in calcium imaging^25^ are not distorting our main findings, we are replicating these experiments using Neuropixels probes. One additional caveat is that the data and the model, as developed so far, are limited to situations where the distribution of contrasts in the environment is independent of stimulus orientation. Relevant scenarios exist where this is not the case. In cases of astigmatism, for example, the contrast of gratings at some orientations is reduced compared to others. Future work will examine how the cortex adapts in such conditions.

## Methods

### Experimental model and subject details

All experimental procedures were approved by UCLA’s Office of Animal Research Oversight (the Institutional Animal Care and Use Committee) and were in accord with guidelines set by the U.S. National Institutes of Health. A total of 9 mice, male (5) and female (4), aged P35-56, were used. These animals were obtained as a cross between TRE-GCaMP6s line G6s2 (Jackson Lab, https://www.jax.org/strain/024742) and CaMKII-tTA (https://www.jax.org/strain/007004). There were no obvious differences in the exponents of the power law between male and female datasets (**Table 1**, Wilcoxon test, *p* > 0.5 for both exponents, *n* = 14 experiments evenly split between male and female), thus we report our data together.

**Table 1.**
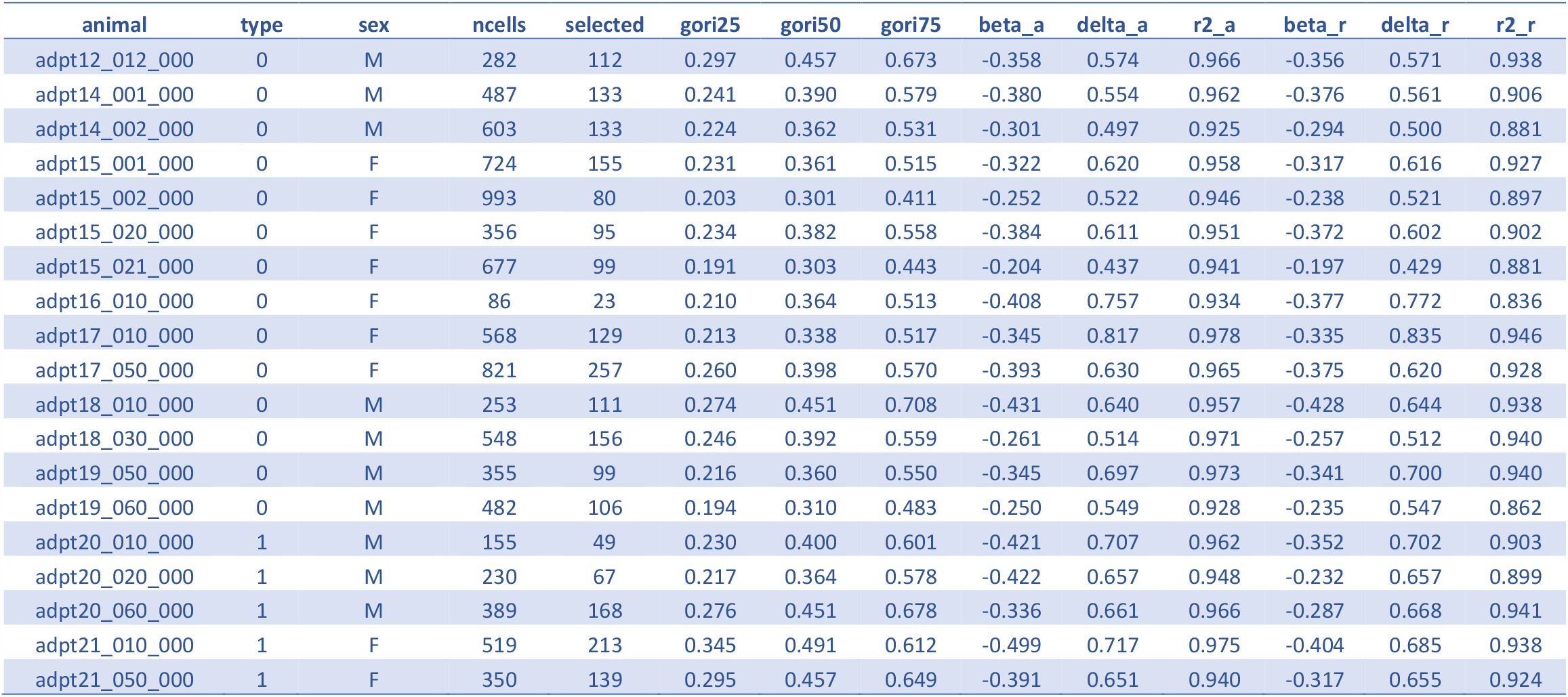
Summary of the datasets. Each row corresponds to an independent experimental session. From left to right, the columns indicate: session name, type of experiment (0 = environments with the same von Mises distribution but different centers, 1 = natural orientation distributions), animal sex, total number of cells segmented, total number of cells that passed the data selection criterion, 25%, 50% and 75% of one minus the circular variance of the responses, exponents estimated via the averaging procedure and the corresponding R-squared value (**Fig 2b**), the exponents estimated by modeling the ratios (**Equation 1**) and the corresponding R-squared values for this model (**Fig 2f**).

| animal | type | sex | ncells | selected | gori25 | gori50 | gori75 | beta_a | delta_a | r2_a | beta_r | delta_r | r2_r |
| --- | --- | --- | --- | --- | --- | --- | --- | --- | --- | --- | --- | --- | --- |
| adpt12_012_000 | 0 | M | 282 | 112 | 0.297 | 0.457 | 0.673 | -0.358 | 0.574 | 0.966 | -0.356 | 0.571 | 0.938 |
| adpt14_001_000 | 0 | M | 487 | 133 | 0.241 | 0.390 | 0.579 | -0.380 | 0.554 | 0.962 | -0.376 | 0.561 | 0.906 |
| adpt14_002_000 | 0 | M | 603 | 133 | 0.224 | 0.362 | 0.531 | -0.301 | 0.497 | 0.925 | -0.294 | 0.500 | 0.881 |
| adpt15_001_000 | 0 | F | 724 | 155 | 0.231 | 0.361 | 0.515 | -0.322 | 0.620 | 0.958 | -0.317 | 0.616 | 0.927 |
| adpt15_002_000 | 0 | F | 993 | 80 | 0.203 | 0.301 | 0.411 | -0.252 | 0.522 | 0.946 | -0.238 | 0.521 | 0.897 |
| adpt15_020_000 | 0 | F | 356 | 95 | 0.234 | 0.382 | 0.558 | -0.384 | 0.611 | 0.951 | -0.372 | 0.602 | 0.902 |
| adpt15_021_000 | 0 | F | 677 | 99 | 0.191 | 0.303 | 0.443 | -0.204 | 0.437 | 0.941 | -0.197 | 0.429 | 0.881 |
| adpt16_010_000 | 0 | F | 86 | 23 | 0.210 | 0.364 | 0.513 | -0.408 | 0.757 | 0.934 | -0.377 | 0.772 | 0.836 |
| adpt17_010_000 | 0 | F | 568 | 129 | 0.213 | 0.338 | 0.517 | -0.345 | 0.817 | 0.978 | -0.335 | 0.835 | 0.946 |
| adpt17_050_000 | 0 | F | 821 | 257 | 0.260 | 0.398 | 0.570 | -0.393 | 0.630 | 0.965 | -0.375 | 0.620 | 0.928 |
| adpt18_010_000 | 0 | M | 253 | 111 | 0.274 | 0.451 | 0.708 | -0.431 | 0.640 | 0.957 | -0.428 | 0.644 | 0.938 |
| adpt18_030_000 | 0 | M | 548 | 156 | 0.246 | 0.392 | 0.559 | -0.261 | 0.514 | 0.971 | -0.257 | 0.512 | 0.940 |
| adpt19_050_000 | 0 | M | 355 | 99 | 0.216 | 0.360 | 0.550 | -0.345 | 0.697 | 0.973 | -0.341 | 0.700 | 0.940 |
| adpt19_060_000 | 0 | M | 482 | 106 | 0.194 | 0.310 | 0.483 | -0.250 | 0.549 | 0.928 | -0.235 | 0.547 | 0.862 |
| adpt20_010_000 | 1 | M | 155 | 49 | 0.230 | 0.400 | 0.601 | -0.421 | 0.707 | 0.962 | -0.352 | 0.702 | 0.903 |
| adpt20_020_000 | 1 | M | 230 | 67 | 0.217 | 0.364 | 0.578 | -0.422 | 0.657 | 0.948 | -0.232 | 0.657 | 0.899 |
| adpt20_060_000 | 1 | M | 389 | 168 | 0.276 | 0.451 | 0.678 | -0.336 | 0.661 | 0.966 | -0.287 | 0.668 | 0.941 |
| adpt21_010_000 | 1 | F | 519 | 213 | 0.345 | 0.491 | 0.612 | -0.499 | 0.717 | 0.975 | -0.404 | 0.685 | 0.938 |
| adpt21_050_000 | 1 | F | 350 | 139 | 0.295 | 0.457 | 0.649 | -0.391 | 0.651 | 0.940 | -0.317 | 0.655 | 0.924 |

### Surgery

Imaging was performed by visualizing activity through chronically implanted cranial windows over primary visual cortex. Carprofen was administered pre-operatively (5mg/kg, 0.2mL after 1:100 dilution). Mice were anesthetized with isoflurane (4%–5% induction; 1.5%–2% surgery). The core body temperature was maintained at 37.5C. The eyes were coated with a thin layer of ophthalmic ointment during the surgery. Anesthetized mice were mounted in a stereotaxic apparatus using blunt ear bars placed in the external auditory meatus. A portion of the scalp overlying the two hemispheres of the cortex was subsequently removed to expose the skull. The skull was dried and covered by a thin layer of Vetbond. After the Vetbond dried (15 min), we affixed an aluminum bracket with dental acrylic. The margins were sealed with Vetbond and dental acrylic to prevent any infections. A craniotomy was performed over monocular V1 on the left hemisphere using a high-speed dental drill. Special care was taken to ensure that the <u>dura</u> was not damaged during the process. Once the skull was removed, a sterile 3 mm diameter cover glass was placed directly on the exposed dura and sealed to the surrounding skull with Vetbond. The remainder of the exposed skull and the margins of the cover glass were sealed with dental acrylic. Mice were allowed to recover on a heating pad and once awake, they were transferred back to their home cage. Carprofen was administered post-operatively for 72 hours. We allowed mice to recover for at least 6 days before the first imaging session.

### Two-photon imaging

We conducted imaging sessions in awake animals starting 6-8 days after surgery. Mice were positioned on a running wheel and head-restrained under a resonant, two-photon microscope (Neurolabware, Los Angeles, CA) controlled by Scanbox acquisition software and electronics (Scanbox, Los Angeles, CA). The light source was a Coherent Chameleon Ultra II laser (Coherent Inc, Santa Clara, CA). Excitation wavelength was set to 920 nm. The objective was an x16 water immersion lens (Nikon, 0.8NA, 3mm working distance). The microscope frame rate was 15.6Hz (512 lines with a resonant mirror at 8kHz). The field of view was 730*μ*m × 445*μ*m. The objective was tilted to be approximately normal the cortical surface. Images were processed using a standard pipeline consisting of image stabilization, cell segmentation and signal extraction using Suite2p (https://suite2p.readthedocs.io/)^26^. A custom deconvolution algorithm was used^27^. We have previously compared the results of different deconvolution algorithms and found little variability in the results^18^. A summary of the experiments, including statistical summaries, are presented in **Table 1**.

### Visual stimulation

We used a Samsung CHG90 monitor positioned 30 cm in front of the animal for visual stimulation. The screen was calibrated using a Spectrascan PR-655 spectro-radiometer (Jadak, Syracuse, NY), generating gamma corrections for the red, green, and blue components via a GeForce RTX 2080 Ti graphics card. Visual stimuli were generated by a custom written Processing 4 sketch using OpenGL shaders (see http://processing.org). At the beginning of each experiment, we obtained a coarse retinotopy map of primary visual cortex as described elsewhere^18^. The center of the aggregate population receptive field was used to center the location of our stimuli in these experiments. Stimuli were presented within a circular window with a radius of 25 deg. The spatial frequency of the gratings was fixed at 0.04 cpd. The orientation domain was discretized in steps of 10 deg, from 0 to 180 deg. The spatial phases at each orientation were uniformly randomized from 0 to 360 deg in steps of 45 deg. Contrast was drawn uniformly form a set 5% × ζ^*q*^ for *q* = 0, …, 6 and ζ = (100/5)^(1/6)^≈ 1.6475. A total of 5400 trials (6 blocks of 5 min each at 3 stim per sec) were collected for each environment. For a uniform environment, this results in an average of 300 trials per orientation. The appearance of a new stimulus on the screen was signaled by a TTL line sampled by the microscope. As a failsafe, we also signaled the onset of the stimulus by flickering a small square at the corner of the screen. The signal of a photodiode was sampled by the microscope as well. In some experiments (**Fig 3c**) the orientation distributions were drawn from natural samples we measured at UCLA as described in a prior study^18^.

### Optimal stimulus-response delay

For each environment, we calculated the magnitude of the population response *T* microscope frames after the onset of the stimulus, where *T* ranged from -2 to 15. The frame rate of the microscope was 15.53 frames/sec. The time to peak of these curves agreed for all environments. We therefore averaged the magnitudes across all the 3 environments and defined the optimal stimulus-response delay as the time (in microscope frames) between the onset of the stimulus and the peak response magnitude of the population (averaged across orientations and contrasts). This calculation was the same for gratings and for movie sequences. When computing the mean population vector for a given stimulus, we also averaged across spatial phases, thus minimizing the effect of eye movements on our analyses.

### Statistics and reproducibility

We conducted experiments by independently measuring the adaptation of neural populations in visual cortex in 19 different instances (see **Table 1**). Linear models were fitted to the data using Matlab’s fitlm() function. The goodness of fit of linear models was evaluated using the *R*^2^ statistic (the coefficient of determination^28^). Both coefficients were highly significant in all experiments (*p*-values less than 10^−6^). As the study did not involve different groups undergoing different treatments, there was no need for randomization or blind assessment of outcomes. Data selection was used to select a subset of neurons with good orientation tuning, attaining a circular variance^29^ of less than 0.5. Cohen’s *d* was computed using Matlab’s meanEffectSize() function for different contrasts levels for a given environment and orientation. We then computed the average discriminability by pooling the data across all orientations and environments (**Fig 3f,g**).

## Data availability

Data including the mean responses of the population for each experiment can be found in the Figshare repository at_______________________________.

## Code availability

Sample code describing the structure of the database and the replication of some of our analyses can be found along with the data at_________________________________.

## Acknowledgements

This study was supported by NS116471 and EY034488 to D. L. R.

## Author contributions

E.T. performed all the animal surgeries. M.D. contributed to manuscript preparation, experimental design, and to the conceptual and theoretical implications of the data. D.L.R. Devised the experiments, wrote the visual stimulus, collected, and analyzed the data, prepared the data for the repository, and wrote the initial version of the manuscript.

